# Positive reward prediction errors strengthen incidental memory encoding

**DOI:** 10.1101/327445

**Authors:** Anthony I. Jang, Matthew R. Nassar, Daniel G. Dillon, Michael J. Frank

## Abstract

The dopamine system is thought to provide a reward prediction error signal that facilitates reinforcement learning and reward-based choice in corticostriatal circuits. While it is believed that similar prediction error signals are also provided to temporal lobe memory systems, the impact of such signals on episodic memory encoding has not been fully characterized. Here we develop an incidental memory paradigm that allows us to 1) estimate the influence of reward prediction errors on the formation of episodic memories, 2) dissociate this influence from other factors such as surprise and uncertainty, 3) test the degree to which this influence depends on temporal correspondence between prediction error and memoranda presentation, and 4) determine the extent to which this influence is consolidation-dependent. We find that when choosing to gamble for potential rewards during a primary decision making task, people encode incidental memoranda more strongly even though they are not aware that their memory will be subsequently probed. Moreover, this strengthened encoding scales with the reward prediction error, and not overall reward, experienced selectively at the time of memoranda presentation (and not before or after). Finally, this strengthened encoding is identifiable within a few minutes and is not substantially enhanced after twenty-four hours, indicating that it is not consolidation-dependent. These results suggest a computationally and temporally specific role for putative dopaminergic reward prediction error signaling in memory formation.

## Introduction

Behaviors are often informed by multiple kinds of memories. For example, a decision about what to eat for lunch might rely on average preferences that have been slowly learned over time and that aggregate over many previous experiences, but it might also be informed by specific, temporally precise memories (e.g., ingredients seen in the fridge on the previous day). These different kinds of memories prioritize different aspects of experience. For example, reinforcement learning typically aggregates information across relevant experience to form general preferences that are used guide behavior ^1^, whereas episodic memories allow access to details about specific, previously experienced events or scenes with limited interference from other similar ones. Computational modeling suggests that these two kinds of memories have different representational requirements and are likely subserved by anatomically distinct brain systems ^2–4^ In particular, a broad array of evidence suggests that reinforcement learning is implemented through cortico-striatal circuitry in the prefrontal cortex and basal ganglia ^5–8^ whereas episodic memory appears to be reliant on synaptic changes in medial temporal structures, especially the hippocampus ^1, 9–12^.

While the distinct anatomy of these systems allows them to operate over different representations, they are not independent. Medial temporal regions provide direct inputs into striato-cortical regions ^2,3,13–15^ and both regions receive shared information through common intermediaries ^5–8,16^. Furthermore, both systems receive neuromodulatory inputs that undergo context dependent fluctuations that can affect synaptic plasticity and alter information processing in both systems ^17,18^ Recent work in computational neuroscience has highlighted potential roles for two neuromodulators, dopamine and norepinephrine, in implementing and optimizing reinforcement learning in a changing world. In particular, dopamine is thought to supply a reward prediction error (RPE) signal that gates Hebbian plasticity in the striatum, facilitating repetition of rewarding actions ^5,6,19,20^. In untrained animals dopamine prediction error signals are observed in response to primary rewards, but with experience dopamine signals precede to the earliest cue predicting future reward ^5^. Such cue-induced dopamine signals are thought to serve a motivational role ^21^, biasing behavior toward the effortful and risky actions that could allow for acquisition of the upcoming reward ^22–27^ In contrast, norepinephrine has been proposed to provide a salience signal that amplifies overall learning from surprising events, irrespective of valence, or during periods of uncertainty ^28–30^. In many experimental tasks, such a salience signal would look very similar to a reward prediction error signal, but careful experimental design can result in their dissociation ^31^.

While normative roles for dopamine and norepinephrine have frequently been discussed in terms of their effects on reinforcement learning and motivational systems, such signals likely also affect processing in medial temporal memory systems ^32–35^. For example, dopamine can enhance LTP ^36^ and replay ^37^ in the hippocampus, which could potentially provide a mechanism to prioritize behaviorally relevant information for longer term storage ^32^. More recent work using optogenetics to perturb hippocampal dopamine inputs revealed a biphasic relationship, whereby low levels of dopamine suppress hippocampal information flow but higher levels of dopamine facilitate it ^35^. Given that the highest levels of dopamine are typically elicited by dopamine bursts ^38^, especially with larger reward prediction errors ^5^, this result suggests that memory encoding in the hippocampus might be enhanced for unexpectedly positive events.

However, despite relatively strong evidence that dopaminergic projections signal reward prediction errors ^5,39^, and that dopamine release in the hippocampus can facilitate memory encoding in non-human animals ^40^, evidence for a positive effect of reward prediction errors on memory formation in humans is scarce. Monetary incentives and reward expectation can be manipulated to improve episodic encoding, even of incidental memories, but it is not clear that such effects are driven by reward prediction errors rather than motivational signals or reward value per se ^18,33,41,42^. The few studies that have closely examined the relationship between reward prediction error signaling and episodic memory have not found evidence for a memory advantage after positive prediction errors ^43,44^. However, there are a number of technical factors that could mask a relationship between reward prediction errors and incidental memory formation in standard paradigms. For instance, such tasks typically have not controlled for salience signals, such as surprise and uncertainty, that can be closely related to RPEs, and that are thought to exert independent effects on episodic encoding through a separate noradrenergic neuromodulatory system ^28–30,45^.

Here we combine a novel behavioral paradigm with computational modeling to clarify the impact of prediction error and surprise, elicited during reinforcement learning, on episodic encoding. Our paradigm required subjects to encode images in the context of a learning and decision-making task, and then to complete a surprise recognition memory test for the images. The decision-making task required subjects to decide whether to accept or reject a risky gamble based on the value of potential payouts and the reward probabilities associated with two image categories, which they learned incrementally based on trial-by-trial feedback. Our design allowed us to measure and manipulate reward prediction errors at multiple timepoints and to dissociate those RPEs from other computational factors that are often correlated with them. In particular, our paradigm and computational models allowed us to manipulate and measure surprise and uncertainty, which have also been implicated in learning and episodic encoding, and are often closely related to RPEs in many tasks. However, both surprise and uncertainty are thought be conveyed through noradrenergic modulation (rather than dopamine, which is thought to reflect RPE). We also assessed the degree to which relationships between these computations and episodic encoding depend on consolidation by testing recognition memories either immediately after the decision task or after a 24 hour delay.

Our results reveal that subjects were more likely to remember images presented in trials in which they accepted the risky gamble. Moreover, the degree of this memory benefit increased with the RPE experienced at the time of image presentation, but not by the RPE, surprise, or uncertainty associated with either the previous or subsequent trial outcome. These results were replicated in an independent sample, which also demonstrated sensitivity to counterfactual information about choices the participants did not make. Collectively, these data demonstrate a key role for reward prediction errors in episodic encoding, clarify the timescale and computational nature of interactions between reinforcement learning and encoding, and make testable predictions about the neuromodulatory mechanisms underlying both processes.

## Results

The goal of this study is to elucidate how computational factors that govern trial-to-trial learning and decision-making might impact episodic memory encoding and retrieval. To do so, we designed a two-part task that included a learning and decision-making phase (Fig 1a) followed by a recognition memory phase (Fig 1c) (see Methods for additional details regarding the task, participants, and analysis). During the learning phase, on each trial subjects decided whether to accept (“play”) or reject (“pass”) an opportunity to gamble based on the potential reward payout. The magnitude of the potential reward was revealed numerically at the start of each trial, but its probability had to be learned via feedback. Subjects were presented with a visual image and were told that the probabilities of reward would depend on whether the (trial-unique) image belonged to one of two categories (animate or inanimate). The reward probability was yoked across categories, such that p(animate) = 1 − p(animate). On each trial, the subject integrated information about the trial payout (selected at random independently for each trial) and learned probability in order to decide whether to “play” or “pass” (Fig 1a). Furthermore, sequential presentation of value, probability and trial outcome information allowed us to manipulate subject reward expectations dynamically within a trial to separately elicit reward prediction errors before, during, and after image presentation (Fig 1b).

The learning phase thus presented subjects with a series of unique stimuli (images from animate/inanimate categories) in a context that allowed for the statistical dissociation of three computational factors thought to govern learning from feedback: prediction error, surprise, and uncertainty. Dissociation of these three factors was achieved in part through the independent manipulation of reward probability and value, and in part through occasional change points in the assignments of the yoked reward probabilities to each of the two image categories (Fig 1d; ^46^). This design yielded estimates of surprise from an ideal observer model that spiked at improbable outcomes, including — but not limited to—those observed after changes in the reward probabilities (Fig 1e). Estimates of uncertainty changed more gradually and tended to be highest during periods following surprise (i.e., when the reward outcomes are volatile, one becomes more uncertain about the learned probability; Fig 1f). Reward prediction errors at time of feedback were highly variable across trials and more related to the probabilistic (and bivalent) trial outcomes than to transitions in the reward structure (Fig 1g).

While repeated-choice bandit tasks typically involve a prediction error at time of feedback, our task also provided some information about the probability of a rewarding outcome during the decision phase, coincident with the image presentation. This allowed us to examine the effects of a separate, image reward prediction error signal (Fig 1b). On some trials, the presented image category was associated with a higher-than-expected reward probability leading to a positive image reward prediction error, whereas on other trials the presented image suggested a lower than expected reward probability leading to a negative image prediction error (Fig 1h). The image prediction errors could be thought of as the dopamine signal that would be expected to occur in response to the probabilistic reward cue provided by the memoranda themselves.

**Figure 1:**
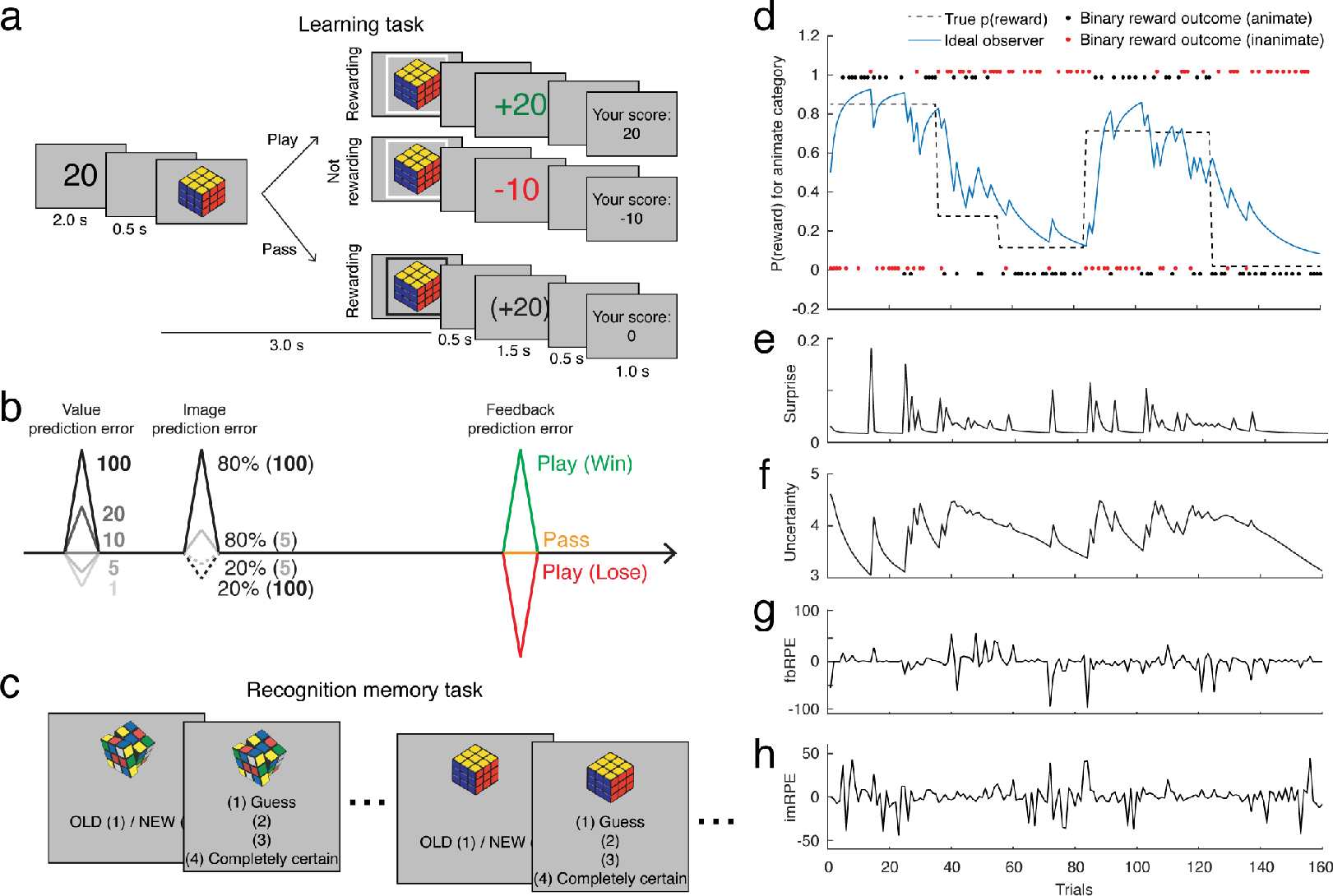
Dissociating effects of reward prediction error, surprise and uncertainty on incidental memory encoding. **a**, In the learning task subjects were informed about the potential value of a successful gamble for the current trial (two seconds, 20 for example trial) and then shown a unique image belonging to one of two categories (animate/inanimate) for a total of three seconds, during which the subject would decide whether to accept (“play”) or reject (“pass”) the gamble. After a brief delay, subjects were informed about their earnings and, if they rejected the gamble, saw counterfactual information regarding the trial outcome. At the end of each trial a cumulative total score was displayed. **b**, The learning task facilitates the manipulation of reward prediction errors before, during, and after image presentation. Prediction errors elicited before image presentation would reflect the value information presented at trial outset (leftmost peak; colors indicate value), prediction errors at image presentation would reflect reward probability information conveyed by the image and its interaction with trial value (middle peak; solid/dashed lines reflect high/low reward probabilities), and prediction errors occurring after image presentation would convey information about trial outcome (rightmost peak; colors reflect trial outcome). **c**, After the learning task, subjects completed a surprise recognition memory test in which each image was either “old” (i.e., from the learning task) or “new” (i.e., a novel lure). Subjects were asked to provide a binary answer along with a 1-4 confidence rating for each image. Importantly, the new images were semantically matched to the old images such that accurate responding depended on the retrieval of detailed perceptual information from encoding. **d**, Reward probabilities during the learning task were determined by image category, were yoked across categories, and were reset occasionally to require learning (black dotted line). Binary outcomes indicating whether the gamble would be rewarded (red/black dots) were governed by reward probabilities and could be used by an ideal observer model to infer the underlying reward probabilities (blue). **e-g**, The ideal observer learned in proportion to the surprise associated with a given trial outcome (**e**) and the uncertainty about its estimate of the current reward probability (**f**), both of which fluctuated dynamically throughout the task and were dissociable from reward prediction error signals at time of feedback (**g**) and at time of image presentation (**h**).

Analyses of data from 199 subjects suggest that they (1) integrated reward probability and value information and (2) utilized reward prediction errors, surprise, and uncertainty to gamble effectively. Subjects increased the proportion of play (gamble) responses as a function of both trial value and the ground truth category reward probability (Fig 2a). To capture trial-to-trial dynamics of subjective category probability assessments, we fit play/pass data from each subject with a set of reinforcement learning models. The simplest such model fit betting behavior as a weighted function of reward magnitude and probability, with probabilities updated on each trial with a fixed learning rate. More complex models (see Methods for details) considered the possibility that this learning rate might itself be adjusted according to other factors such as surprise, uncertainty, or whether the subject had decided to gamble on the trial. Consistent with previous work ^30,31,47,48^, the best fitting model adjusted learning according to normative measures of both surprise and uncertainty (Fig 2c). Coefficients describing the effects of surprise and uncertainty on learning tended to be positive across subjects, indicating that subjects were more responsive to feedback if it was surprising or was provided during a period of uncertainty (Fig 2d; surprise: t = 2.69, df = 198, p = 0.0078; uncertainty t = 6.38, df = 198, p < 0.001). Thus, subject behavior in the learning task is best described as using surprise and uncertainty to scale the extent to which reward prediction errors are used to adjust subsequent behavior.

**Figure 2:**
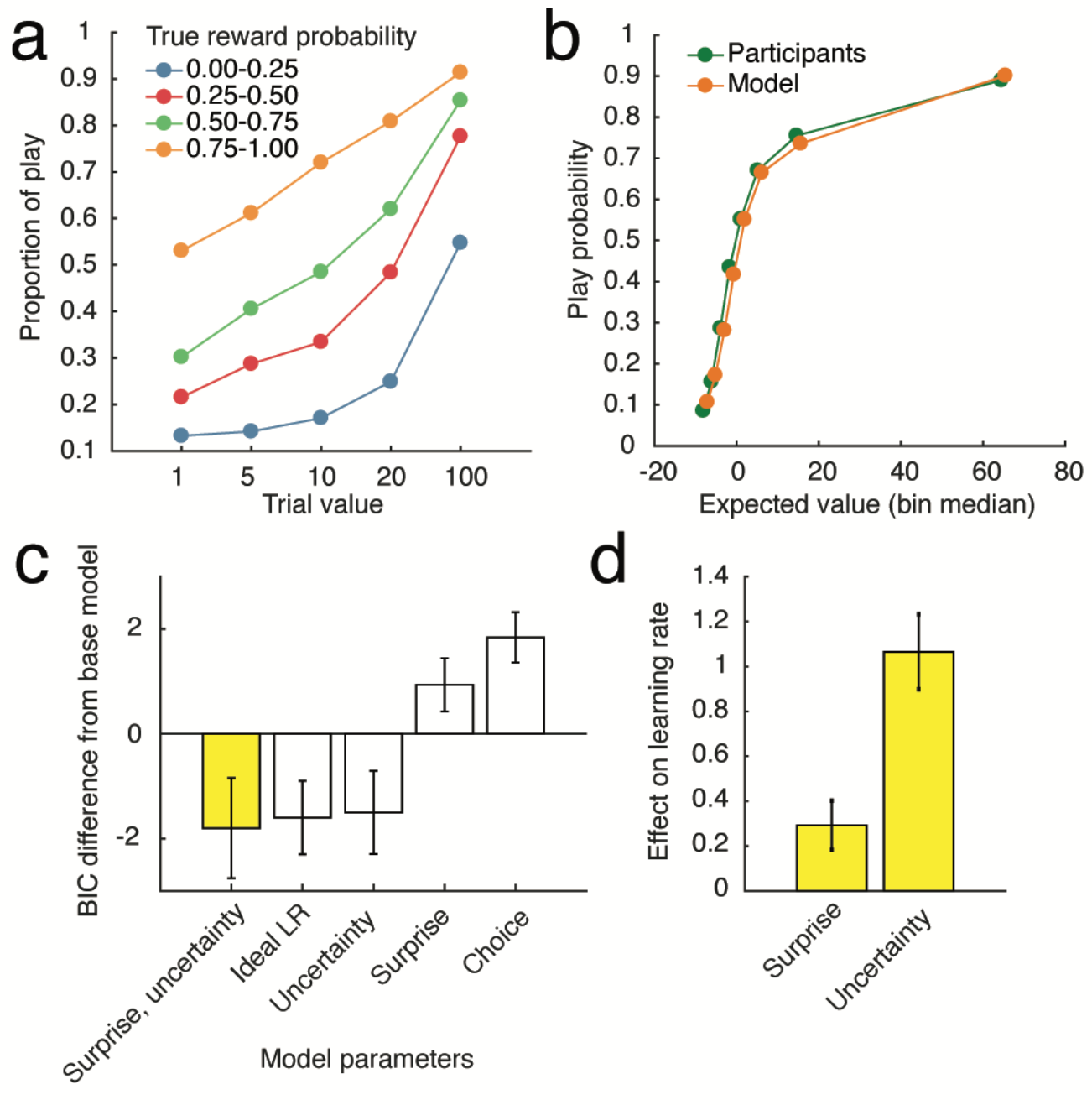
Subjects’ gambling behavior integrated reward value and subjective reward probability estimates, which were updated as a function of surprise and uncertainty. **a**, Proportion of trials in which the subjects chose to play, broken down by reward value and true reward probability. **b**, Subject choice behavior and model-predicted choice behavior. The model with the lowest BIC, which incorporated the effects of surprise and uncertainty on learning rate, was used to generate model behavior. Expected values for all trials were divided into 8 equally sized bins for both subject and model-predicted behavior. **c**, Bayesian information criterion of five reinforcement learning models with different parameters that affect learning rate. **d**, Mean maximum likelihood estimates of surprise and uncertainty parameters of the best fitting model (first bar on **c**). Error bars indicate standard error across subjects.

Behavior in the recognition task confirmed that subjects formed memories about the unique images displayed in the learning task. Subjects completed a surprise recognition memory task either five minutes (no delay, *n* = 109) or twenty-four hours (24 hour delay, *n* = 90) after completion of the learning task. During the recognition memory task, subjects viewed all the “old” images from the learning task plus a semantically matched set of “new” foil images that were not shown previously. For each image, subjects provided a binary response indicating whether the image was old or new, along with a 1-4 confidence measure.

Subjects in both delay conditions reliably identified images that had been presented in the learning task with accuracy above chance (Fig 3a; mean(sem) d’ = 0.85(0.042) for no delay and 0.50(0.030) for 24 hour delay condition). The reliability of memory reports was greater when subjects expressed higher levels of confidence (mean(sem) d’ = 1.12(0.062) for no delay and 0.65(0.039) for 24 hour delay condition) than when they reported lower confidence (mean(sem) d’ = 0.44(0.038) for no delay and 0.25(0.037) for 24 hour delay condition).

To aggregate information provided in the binary reports and confidence measure we transformed these sources of data to create a single 1-8 memory score, such that 8 reflected a high confidence “old” response and 1 reflected a high confidence “new” response. As expected, the true proportion of “old” images was higher when memory scores were highest, and the relationship between memory scores and ground truth was monotonic and roughly linear across both delay conditions (Fig 3b). Thus, subjects incidentally formed lasting memories for the images displayed in the learning task, and memory scores provided a measure of the subjective memory strength associated with each image.

Recognition memory for an image depended on the context in which that image was presented during the learning task. Memory scores for previously viewed images were higher for images observed on trials in which subjects gambled (play) than for images observed on trials in which they passed (Fig 3c; Fig S1). The difference between memory scores for old and new items was larger for play than pass trials (t=3.30, dof=198, p =0.001) and this did not differ across delay conditions (t=0.38, dof=198, p = 0.70) (Fig 3d). Larger memory scores were produced, at least in part, by an increase in the sensitivity of memory reports. Across all possible memory scores, hit rate was higher for play trials than for pass trials and the area under subject-specific ROC curves generated in this way was greater for play as compared to pass trials (Fig 3e; t=3.12, dof=198, p = 0.002) but did not differ across delay conditions (t=0.34, dof=197, p=.73). In principle, this enhanced memory encoding could be driven by positive reward prediction errors, which would occur in response to high probability images (figure 1b) and motivate play decisions (figure 2a). In subsequent analyses we test this idea directly.

**Figure 3:**
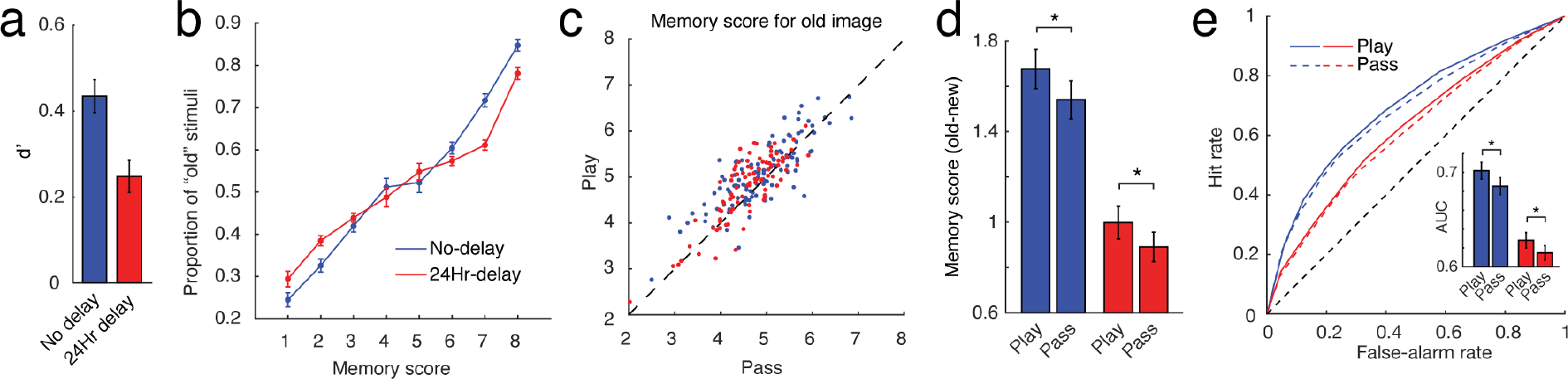
Recognition memory is stronger for stimuli presented on trials in which subjects accepted the gamble. **a**, Average d’ for the two delay conditions. **b**, Average proportion of image stimuli that were “old” (presented during the learning task), separated by memory score. **c**, Mean memory score of “old” images for play trials versus pass trials. Each point represents a unique subject. A majority of subjects lie above the diagonal, indicating better memory performance for play trials. **d**, Mean pairwise difference in memory score between the “old” images and their semantically-matched foil images. **e**, ROC curves for play vs. pass trials. Area under the ROC curves (AUC) shown in the inset. AUC was greater for play versus pass trials, indicating better detection of old versus new images for play trials compared to pass trials. Colors indicate time between encoding and memory testing; blue = no delay, red = 24 hour delay.

The degree to which memory was enhanced on play trials depended on the magnitude of the reward prediction error at time of image presentation (see Fig 1a,b). In particular, memory scores for play trials increased as a function of the reward prediction error computed at the time of image presentation (Fig 4a,b; t = 4.29, df = 198, p = 3 × 10^−5^) in a manner that did not depend on delay condition (t = 0.09, df = 198, p = 0.93). Moreover, this effect was most prominent in the subjects that displayed gambling behaviors that were the most sensitive to trial-to-trial fluctuations in probability and value (spearman correlation of gambling GLM coefficients (probability & value) with memory score image RPE coefficient: ρ = 0.14, p = 0.04).

To better understand the nature of this relationship, we show subsequent memory scores related to the constituent components of the reward prediction error signal. A key feature of an error in reward prediction at the time of image presentation is that it should depend directly on reward probability (i.e., the probability associated with the image category relative to the average reward probability across categories) as this is the reward information cued by the image. In contrast, the trial value should not directly influence RPE because this value was already indicated prior to image presentation, and should thus only inform reward predictions themselves, and not their errors. Consistent with a selective effect of RPE at time of image presentation, subsequent memories were stronger for play trials in which the image category was associated with a higher reward probability (Fig 4c,d; t = 3.31, df = 198, p = 0.001) but not for play trials with higher potential outcome value, which if anything were associated with slightly lower memory scores (Fig 4e,f; t = −1.97, df = 198, p = 0.051). Reward probability effects were stronger in subjects that displayed more sensitivity to probability and value in the gambling task (spearman correlation of gambling GLM coefficients (probability & value) with memory score image reward probability coefficient: ρ = 0.16, p = 0.02).

**Figure 4:**
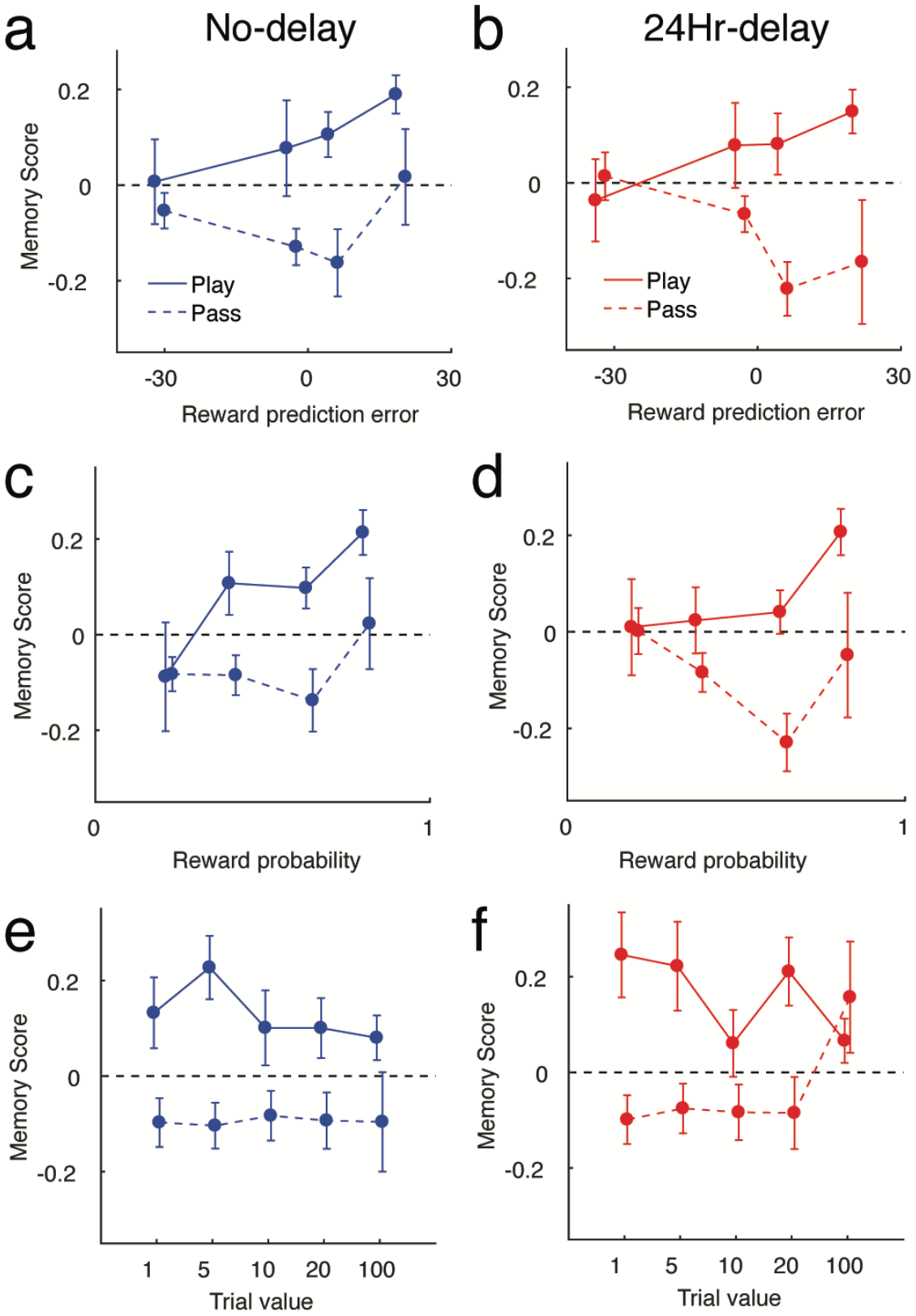
Subsequent memory strength depends on the reward prediction error at time of image presentation, but not trial value. **a,b,** Reward prediction error during image presentation shows a positive association with subsequent recognition memory for the image in both no delay (**a**; blue) and 24 hour delay (**b**; red) conditions. **c-f**, Reward probability estimates (**c,d**), but not reward value (**e,f**) retained the positive association with subsequent memory. This suggests that prediction error that occurs during image presentation, but not the overall value of the image, is driving the subsequent memory effect. Colors indicate time between encoding and memory testing; blue = no delay, red = 24 hour delay.

While subject gambling behavior depended critically on the reward prediction error, uncertainty, and surprise associated with trial feedback, to our surprise none of these factors influenced subsequent memory for the images. Specifically, memory scores were not systematically related to the reward prediction error experienced at time of feedback on the trial preceding image presentation (Fig 5a; t = 0.25, dof = 198, p = 0.80) or immediately after image presentation (Fig 5b; t = −1.19, dof = 198, p = 0.26). Similarly, the surprise associated with feedback preceding (Fig 5b; t = −1.71, dof = 198, p = 0.088) or following (Fig 5d; t = 1.24, dof = 198, p = 0.16) image presentation was not systematically related to subsequent memory scores. Furthermore, there were no obvious relationships between subsequent memory score and uncertainty, again despite the apparent impact of uncertainty on subject gambling behavior (fig 5e, previous trial uncertainty, t=1.70, dof=198, p=0.091; fig 5f, current trial uncertainty, t = −0.63, dof = 198, p = 0.53).

**Figure 5:**
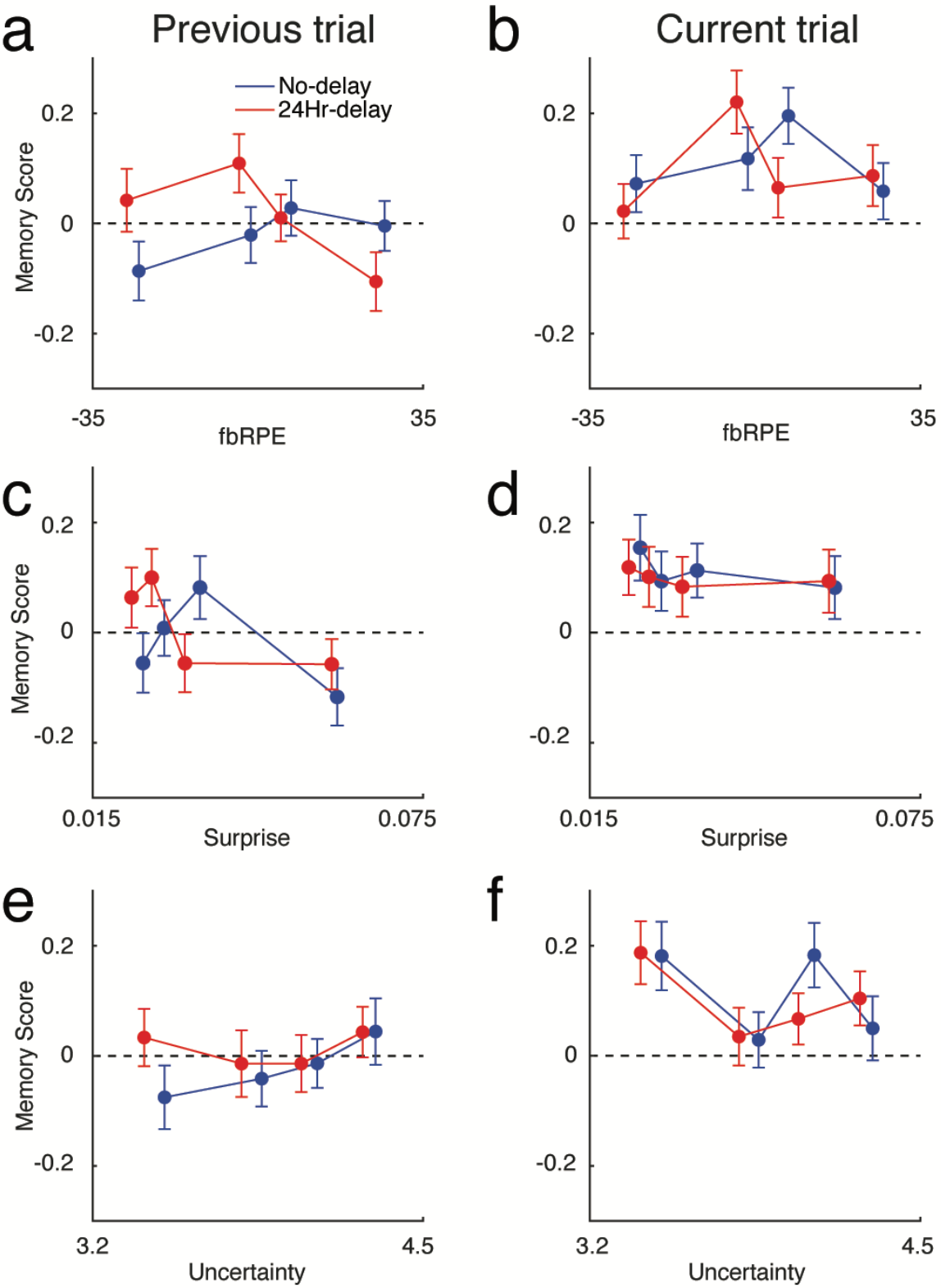
Subsequent memory is not affected by unexpected rewards, surprise, or outcome uncertainty during the feedback preceding or following image presentation. **a,b,** Reward prediction error during the feedback phase of the previous trial (**a**) or current trial (**b**) did not affect subsequent memory. **c,d,** Surprise associated with the feedback phase of the previous (**c**) or current (**d**) trial did not affect subsequent memory. **e,f,** Uncertainty during the previous (**e**) or current (**f**) trial did not affect subsequent memory. Colors indicate time between encoding and memory testing; blue = no delay, red = 24 hour delay.

To better estimate the contributions of learning-related computations to subsequent memory strength, we constructed a hierarchical regression model capable of 1) pooling information across subjects and delay conditions in an appropriate manner, 2) estimating the independent contributions of each factor while simultaneously accounting for all others, and 3) accounting for the differences in memory scores attributable to the images themselves. The hierarchical regression model attempted to predict memory scores by estimating coefficients at the level of items and subjects, as well as estimating the mean parameter value over subjects and the effect of delay condition for each parameter (Fig 6a).

Consistent with the results presented thus far, the hierarchical regression results support the notion that encoding was strengthened by the decision to gamble (play vs. pass) and reward prediction errors elicited at the time of image presentation, but not by the computational factors that controlled learning rate (surprise and uncertainty). Play trials were estimated to contribute positively to encoding, as indexed by uniformly positive values for the posterior density on the play/pass parameter (Fig 6b top row of column 2; table 1). The reward probability associated with the displayed category was positively related to subsequent memory on play trials (Fig 6b column 3; table 1), as was its interaction with value (Fig. 6b column 5; table 1)–although there was no reliable effect of value itself (Fig 6b columns 6; table 1). The direction of the interaction effect suggests that subjects were more sensitive to image probability on trials in which there were more points available to be won, consistent with the more nuanced predictions of a reward prediction error at time of image presentation (Fig 1b). All observed effects were selective for the old items that subjects observed in the task, as the same model fit to the new, foil images yielded coefficients near zero for each of these terms (Fig S2). Consistent with our previous analysis, coefficients for the uncertainty and surprise terms were estimated to be near zero (Fig 6b rightmost columns; table 1).

In addition to supporting our previous analyses, our model allowed us to examine the extent to which any subsequent memory effects required a consolidation period. In particular, any effects on subsequent memory that were stronger in the 24hr delay condition vs. the immediate condition might reflect an effect of post-encoding processes. Despite evidence from animal literature that dopamine can robustly affect memory consolidation (e.g., Bethus et al., 2010), we did not find strong support for any of our effects being consolidation dependent (note lack of positive coefficients in bottom row of Fig 6b, which would indicate effects stronger in the 24 hour condition). As might be expected, subjects in the no delay condition tended to have higher memory scores overall (Fig 6b bottom of column 1; table 1) but their memory scores also tended to change more as a function of reward probability (Fig 6b bottom of column 3; table 1) than did their counterparts in the 24hr delay condition. These results reveal the expected decay of memory over time, and suggest that the image prediction errors induced by categories associated with higher reward probability are associated with an immediate and decaying boost in memory accuracy.

In summary, behavioral data and computational modeling support a role for computation of surprise, uncertainty and reward prediction error in the learning phase of our paradigm. However, only decisions to gamble and the instantaneous reward prediction error at the time of image presentation were related to subsequent memory. To better understand the reward prediction error effect, and to ensure the reliability of our findings, we conducted a second experiment.

**Figure 6:**
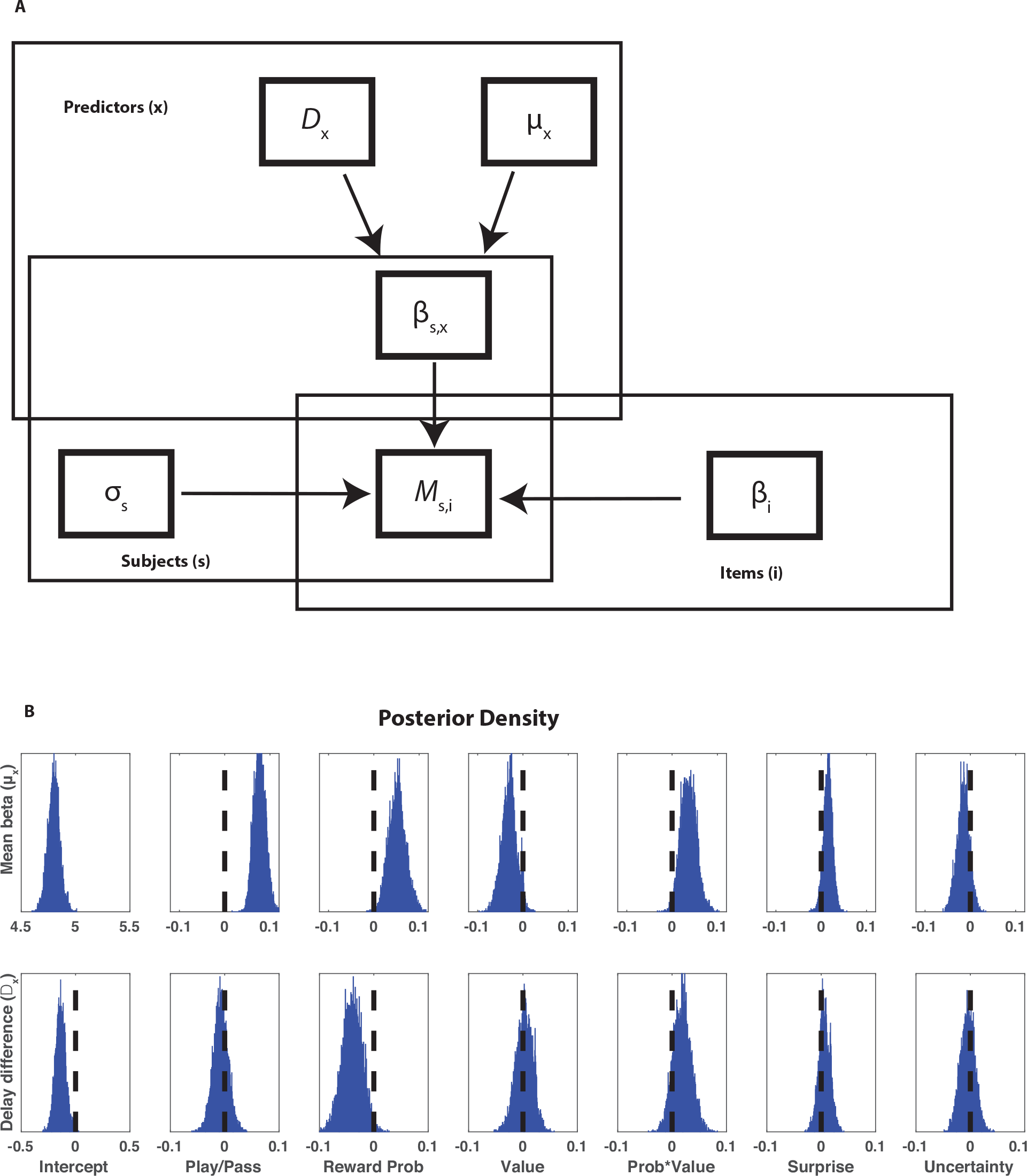
Hierarchical regression model reveals effects of choice and positive prediction errors on recognition memory encoding. **a:** Graphical depiction of hierarchical regression model. Memory scores for each subject and item (*M*_s,i_) were modeled as normally distributed with subject specific variance (σ_s_) and a mean that depended on the sum of two factors: 1) subject level predictors related to the decision context in which an image was encountered (i.e., whether the subject played or passed) linearly weighted according to coefficients (β_s,x_) and 2) item level predictors specifying which image was shown on each trial and weighted according to their overall memorability across subjects (β_i_). Coefficients for subject level predictors were assumed to be drawn from a global mean value for each coefficient (μ_x_) plus an offset related to the delay condition (*D*_x_). Parameters were weakly constrained with priors that favored mean coefficient values near zero and low variance across subject and item specific coefficients. **b:** Posterior probability densities for mean predictor coefficients (μ_x_; top row) and delay condition parameter difference (*D*_x_; bottom row) estimated through MCMC sampling over the graphical model informed by the observable data (*M*_s,i_).

## Experiment 2

Our previous findings suggested that variability in the strength of memory encoding was related to computationally-derived RPE signals and the gambling behavior that elicited them. However, the yoked reward probabilities in Experiment 1 ensured that the reward probability associated with the image category presented would be perfectly anti-correlated with the reward probability associated with the other category that was not presented. Thus, while a high reward category item would increase the expected reward relative to before the trial and hence elicit an RPE, we were unable to disentangle whether the observed effects were driven by the reward probability directly, the counterfactual reward associated with the alternate category, or, as would be predicted by a true reward prediction error, their difference. To address this issue we conducted a second experiment in which expectations about reward probability were manipulated independently of the actual reward probability on each trial allowing us to distinguish between these alternative explanations.

Specifically, we modified the design of our task such that the learning phase included separate manipulations of reward probability for the two image categories. Thus, during some trials both categories would be associated with a high reward probability and in some sessions both would be associated with a low reward probability (Fig 7a-b). In this design, RPEs are relatively small when the reward probabilities are high for both categories and deviate much more substantially when the reward probability differs across image category (Fig 7c). Thus, if the factor boosting subsequent memory scores is truly a reward prediction error, it should depend positively on the reward probability associated with the image category, but negatively with the reward probability associated with the other category.

Subjects in both delay conditions reliably identified images presented in the learning task with accuracy above chance (Fig 7d; mean(sem) d’ = 0.91(0.058) for no delay and 0.54(0.034) for 24 hour delay condition). We also observed a robust replication of the difference in memory scores for trials in which subjects either gambled or passed on the offer to gamble (Fig 7e; t=3.89, dof=172, p<0.001; difference across delay condition, t=−0.60, dof=171, p=0.55).

In the new experimental design, we could analyze variability in memory scores for each old image as a function of its associated category reward probability (“image category”) and the reward probability associated with the other category (“other category”). Based on the RPE account, we hypothesized that we would see a positive effect of the experienced reward (image category) and a negative effect of the reward that would have otherwise been available (other category) on subsequent memory. Consistent with this prediction, for both delay conditions, there was a cross-over effect whereby memory scores scaled positively with the probability associated with the image category (Fig 7f,g t=2.37, dof=172, p=0.019), and negatively with the probability associated with the other category (t=−2,27, dof=172, p=0.024). These effects did not differ across delay conditions (current category, t=0.55, dof=171, p=0.58; other category, t=0.36, dof=171, p=0.72).

**Figure 7:**
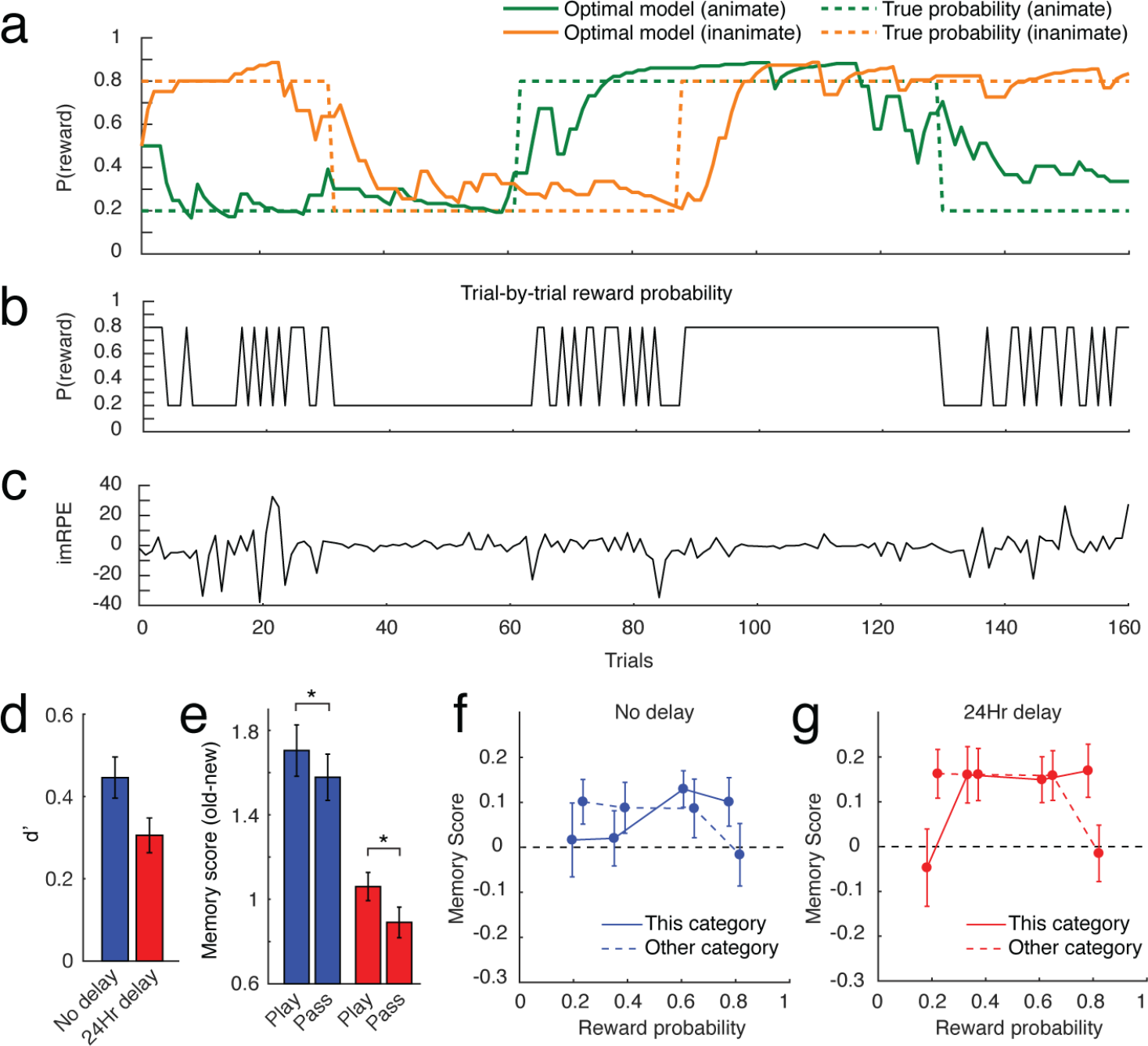
Experiment 2 allowed us to separately estimate the effects of both category probabilities on subsequent memory recall. **a:** In the new learning task, the true reward probabilities of the two categories were independent, and were restricted to either 0.2 or 0.8. The change-points occurred randomly, ensuring that each subject completed a unique learning task. Restrictions were applied so that each task contained at least one block (constituting at least 20 trials) of the four possible reward probability combinations (0.2/0.2, 0.2/0.8, 0.8/0.2, 0.8/0.8). **b:** Trial-by-trial reward probability shows stretches of stable reward probability (0.2/0.2, 0.8/0.8), or varying reward probability (0.2/0.8, 0.8/0.2). **c;** The variability of imRPE is influenced by the reward probability conditions. **d:** Average d’ for both delay conditions. **e**: Mean pairwise difference in memory score between the “old” images and their semantically-matched foil images. **f,g**: There is a positive association between reward probability of the currently shown stimulus category and subsequent recognition memory, and a negative association between the reward probability of the other stimulus category and memory. In panels d-f, colors indicate time between encoding and memory testing; blue = no-delay, red = 24 delay.

To better estimate the effects of image category, other category, and play/pass behavior on subsequent memory we fit memory score data with a modified version of the hierarchical regression model that included separate reward probability terms for the “image” and “other” categories. Posterior density estimates for the play/pass coefficient were greater than zero (Fig 8a; table 1), replicating our previous finding. The posterior density for the “image category” and “other category” probabilities was concentrated in the region over which image category was greater than other category (mean [95% CI] image category coefficient – other category coefficient: 0.052 [0.015,0.94]) and supported independent and opposite contributions of both category probabilities (Fig 8b; table 1). These results, in particular the negative effect of “other category” probability on the subsequent memory scores, are more consistent with an effect of prediction error than a direct effect of reward prediction itself. More generally, these results support the hypothesis that reward prediction errors elicited at the time of image presentation enhance the degree to which such images are encoded in episodic memory systems.

**Figure 8:**
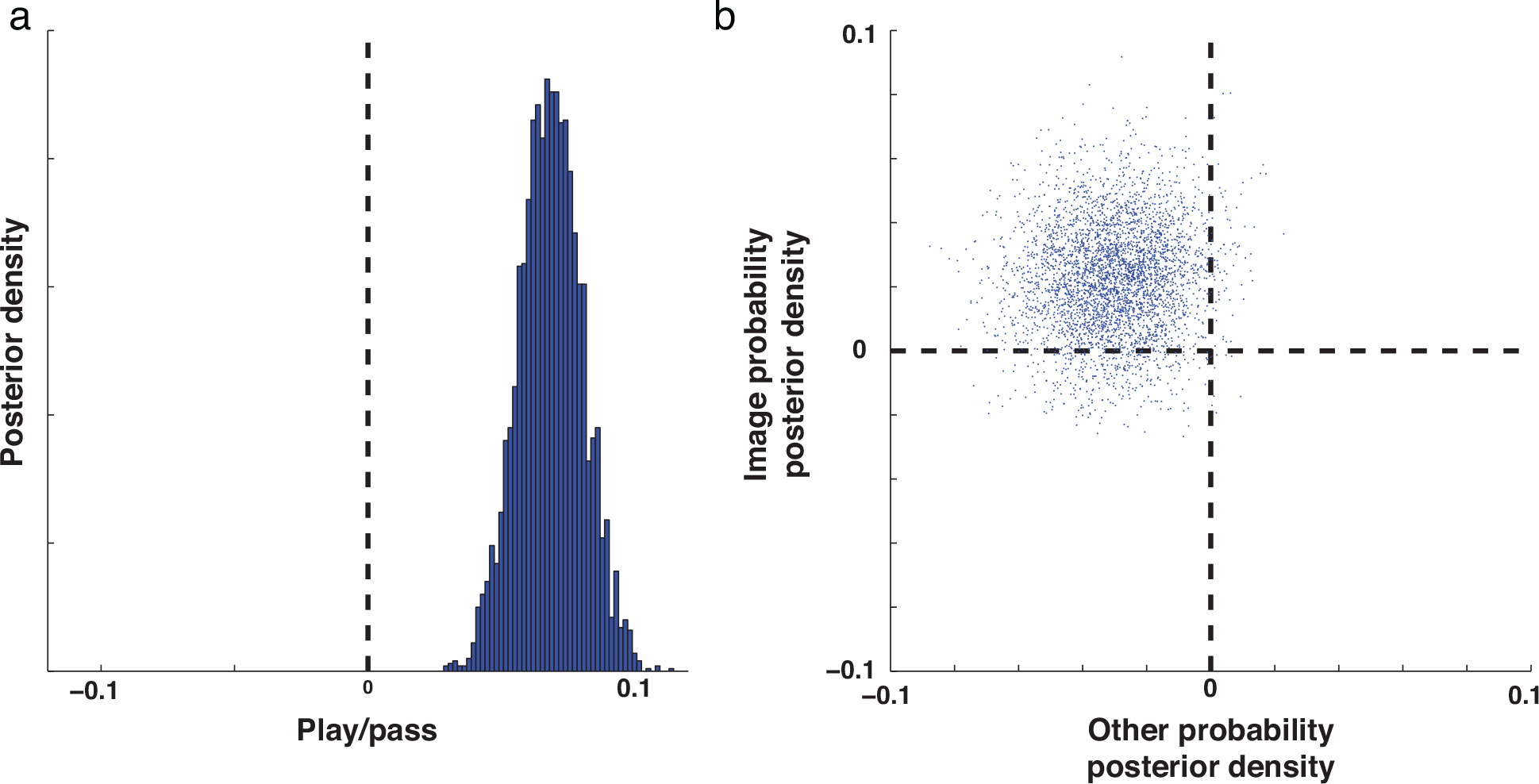
Memory scores in experiment two depend on subject gambling behavior and the probabilities associated with both image categories. Memory score data from experiment two was fit with a version of the hierarchical regression model described in figure 6A to replicate previous findings and determine whether reward probability effects were attributable to both observed and unobserved category probabilities. **a:** Posterior probability estimates of the mean play/pass coefficient were greater than zero and consistent with those measured in the first experiment. **b:** Image category probability (observed) coefficients are plotted against other category probability (unobserved) coefficients revealing that subjects tended to have higher memory scores for images that were associated with high reward probabilities (upward shift of density relative to zero) and when the unobserved image category was associated with a low reward probability (leftward shift of density relative to zero).

Despite the general agreement between the two experiments, there was one noteworthy discrepancy. While hierarchical models fit to both datasets indicated higher probability of positive coefficients for the interaction between value and probability (e.g. positive effects of probability on subsequent memory are greater for high value trials), the 95% credible intervals for these estimates in experiment two included zero as a possible coefficient value (table 1) indicating that the initial finding was not replicated in the strictest sense.

To better understand this discrepancy, and make the best use of the data from both experiments we extended the hierarchical regression approach to include additional coefficients capable explaining differences across the two experiments and fit this extended model to the combined data. As expected, this model provided evidence for a memory advantage on play trials, and an amplification of this advantage for trials with a high reward probability based on image category (Fig S4; table 1). Across the combined dataset there was also positive effect of the interaction between value and probability (Fig S4; table 1), supporting our initial observation in experiment one. Furthermore, we observed that the reward probability effect was greater in the no delay condition (Fig S4; table 1), with no evidence for any memory effects being stronger in the 24 hour delay condition (table 1; all other delay difference ps >.19).

## Discussion

An extensive prior literature has linked dopamine to reward prediction errors elicited during reinforcement learning ^5,6,19,39,49–51^, and a much smaller literature has suggested that dopamine can also influence the encoding and consolidation of episodic memories by modulating activity in the medial temporal lobes ^18,40,52^. To date, however, there has been mixed evidence regarding the relationship between prediction error signaling and memory encoding. Here we used a novel two-stage learning and memory paradigm along with computational modeling to better characterize how prediction error signals affect the strength of incidental memory formation.

We found that memory encoding was stronger for trials in which the subjects observed an image that was associated with high reward probability (figure 5c,d). This effect was only evident for trials in which subjects accepted the risky offer (consistent with the fact that trial outcome was always zero on pass trials), was evident even after controlling for other potential confounds (figure 6b column 3), and was amplified for trials in which more points were on the line (figure 6b column 5). These results are all consistent with a direct effect of reward prediction error at time of image presentation on memory encoding (figure 5a,b). This interpretation is bolstered by evidence that individuals that were more sensitive to value and probability in the decision making task showed reward prediction error memory benefits to a greater degree. Experiment two further supported the reward prediction error interpretation by demonstrating that memory benefits were composed of equal and opposite contributions of the reward probability associated with the observed image category and that of the unobserved, counterfactual, one (Figure 7f,g, Figure 8b). Together, these results provide evidence for the hypothesis that reward prediction errors enhance the encoding of simultaneously presented incidental visual information for subsequent memory.

We also found that subjects encoded memoranda to a greater degree on trials in which they selected a risky bet (figure 3). This finding is consistent with a positive relationship between prediction error signaling and memory strength, in that subject behavior provides a proxy for the subjective reward probability estimates (figure 2A). However, this behavioral effect was prominent in both experiments, even after controlling for model-based estimates of reward prediction error (figures 6b & 8a). Therefore, while we suspect that this result may at least partially reflect the direct impact of reward prediction error, it may also reflect other factors associated with risky decisions. On play trials, subjects view items while anticipating the uncertain gain or loss of points during the upcoming feedback presentation whereas on pass trials, subjects are assured to maintain their current score. The possibility that this difference in risk might contribute to the subsequent memory effects observed for choice behavior would be consistent with recent work showing that memoranda presented immediately prior to feedback are better remembered if they preceded more uncertain feedback ^53^. One potential confound for these choice effects is the heightened state of attention that might occur before receiving a more informative task outcome. While we are unable to rule this possibility out completely, our study minimizes this possibility by presenting counterfactual information on pass trials that is nearly identical to the experienced outcome information. We find that subjects are slightly more influenced by outcome information provided on play trials, suggesting that anticipatory attention might differ somewhat between the two conditions (Fig S3a); however the degree of this difference was small enough that model selection favored a model that did not distinguish between play and pass trials for learning (Fig 2c). Furthermore, there was no relationship between the degree to which subjects modulated learning from feedback on play versus pass trials and the degree to which they showed subsequent memory improvements on play trials (Fig S3b). Given that risky decisions tend to be preceded by higher levels of dopamine, the possibility of a direct effect of risk taking on subsequent memory would still be consistent with a dopaminergic mechanism ^22–24,27^.

This relationship between reward prediction errors and memory is consistent with a broad literature highlighting the effects of dopaminergic signaling on hippocampal plasticity ^32,35,36^ and memory formation ^40^ as well as an equally broad literature suggesting that dopamine provides a reward prediction error signal ^5,19^ through projections that extend both to the striatum and the hippocampus ^17^. Our results support the behavioral consequences that might be predicted to result from such mechanisms, however they also refine them substantially. In particular, we show that the timing of reward prediction error signaling relative to the memorandum is key; we saw no effect of the reward prediction error elicited by prior or subsequent feedback on memory strength (Figure 4a&b), despite strong evidence that this feedback was used to guide reinforcement learning and decision making (Figure 2). Furthermore, we showed that prediction error effects on memory emerged immediately after task performance and were not enhanced after 24 hours (Figure 5a&b; Figure 6b column 3). These results are somewhat at odds with previous literature suggesting that dopamine dependent memory enhancement emerges only after an extended consolidation period ^40^. It is unclear to what extent we should expect generalization of these results to our study, given the differences in experimental paradigm, timescale, memory demands, and species in the two paradigms. However, our results open the door for future research to (1) directly test whether prediction error driven memory enhancements are mediated by changes in dopamine, and (2) characterize the conditions under which dopamine mediated changes to memory encoding do and do not require a consolidation period.

More generally, our results provide insight into the apparent inconsistency in previous behavioral studies that have attempted to link reward prediction error signals to memory encoding. Consistent with previous work (e.g. ^54^), our results emphasize the importance of choice in the degree to which image valence contributed to memorability. Indeed, for trials in which the subjects passively observed outcomes, we saw no relationship between model derived reward prediction error estimates and subsequent memory strength (Figure 5a&b dotted lines). This might help to explain the lack of a signed relationship between reward prediction errors and subsequent memory strength in a recent study by Rouhani and colleagues that leveraged a Pavlovian design that did not require explicit choices to be made ^44^. In contrast to our results, Rouhani and colleagues observed a positive effect of absolute prediction error, similar to our model-based surprise estimates, on subsequent memory. While we saw no effect of surprise on subsequent memory, other work has highlighted a role for such signals as enhancing hippocampal activation and memory encoding ^55,56^. One potential explanation for this discrepancy is in the timing of image presentation. Our study presented images only briefly during the choice phase of the decision task. By contrast, Rouhani and colleagues presented the memoranda for an extended period that also encompassed the epoch containing trial feedback, potentially explaining why they observed effects related to outcome surprise. More generally, the temporally selective effects of reward prediction error observed here suggest that the reward prediction error effects may differ considerably from other, longer timescale manipulations thought to enhance memory consolidation through dopaminergic mechanisms ^18,33,41,42^

Our results appear somewhat incompatible on first glance with those of Wimmer and colleagues ^43^, who show that stronger prediction error encoding in the ventral striatum is associated with weaker encoding of incidental information. We suspect that the discrepancy between these results is driven by differences in the degree to which memoranda are task relevant in the two paradigms. In our task, subjects were required to encode the memoranda sufficiently to categorize them in order to perform the primary decision-making task, whereas in the Wimmer study, memoranda were unrelated to the decision task and thus might not be well-attended on all trials. Taken together, these results suggest that reward prediction errors are most likely to enhance memory when they are elicited by the memoranda themselves, with the potential influence of secondary tasks eliminated or at least tightly controlled.

In summary, our results demonstrate a role for reward prediction errors in prioritizing information for memory storage. We show that this role is temporally and computationally precise, independent of consolidation duration (at least in the current paradigm), and contingent on decision-making behavior. These data should help clarify inconsistencies in the literature regarding the relationship between reward learning and memory, and they make detailed predictions for future studies exploring the relationship between dopamine signaling and memory formation.

## Methods

### Experiment 1

#### Experimental procedure

The task consisted of two parts: the learning task and memory task. The learning task was a reinforcement learning task with random change-points in reward contingencies of the targets. The memory task was a surprise recognition memory task using image stimuli that were presented during the learning task and foils.

Subjects completed either the no delay or 24-hour delay versions of the task. In the no delay condition, the memory task followed the learning task only after a short break, during which a demographic survey was given. Therefore, the entire task was performed in one sitting. In the 24-hour delay condition, subjects returned 20-30 hours after completing the learning task to do the memory task.

#### Subjects

A total of 287 subjects (142, no delay condition; 145, 24hr-delay condition) completed the task via Amazon Mechanical Turk. From this, 88 subjects (33, no delay; 55, 24hr-delay) were excluded from analysis because they previously completed a prior version of the task or didn’t meet our criteria of above-chance performance in the learning task. To determine whether a subject’s performance was above-chance, we simulated random choices using the same task structure, then computed the total score achieved by the random performance. We then repeated such simulations 5000 times, and assessed whether the subject’s score was greater than 5% of the score distribution from the simulations. The final sample had a total of 199 subjects (109, no delay, 90, 24hr-delay; 101 males, 98 females) with the age of 32.2 ± 8.5 (mean ± SD). Informed consent was obtained in a manner approved by the Brown University Institutional Review Board.

#### Learning task

The learning task consisted of 160 trials, where each trial consisted of three phases – value, image, and feedback (Figure 1a). During the value phase, the amount of reward associated with the current trial was presented in the middle of the screen for 1.5 s. This value was equally sampled from [1, 5, 10, 20, 100]. After an interstimulus interval (ISI) of 0.5 s, an image appeared in the middle of the screen for 3 s (image phase). During the image phase, the subject made one of two possible responses using the keyboard: PLAY (press 1) or PASS (press 0). When a response is made, a colored box indicating the subject’s choice (e.g. black = play, white = pass) appeared around the image. The pairing of box color and subject choice was pseudorandomized across subjects. After this image phase, an ISI of 0.5 s followed, after which the trial’s feedback was shown (feedback phase).

Each trial had an assigned reward probability, such that if the subject chose PLAY, they would be rewarded according to that probability. If the subject chose PLAY and the trial was rewarding, they were rewarded by the amount shown during the value phase (Figure 1a). If the subject chose PLAY but the trial was not rewarding, they lost 10 points regardless of the value of the trial. If the choice was PASS, the subject neither earned nor lost points (+0), and was shown the hypothetical result of choosing PLAY (Figure 1a). During the feedback phase, the reward feedback (+value, −10, or hypothetical result) was shown for 1.5s, followed by an ISI (0.5 s), and a 1 s presentation of the subject’s total accumulated score.

All image stimuli belonged to one of two categories: animate (e.g. whale, camel) and inanimate (e.g. desk, shoe). Each image belonged to a unique exemplar, such that there were no two images of the same animal or object. Images of the two categories had reward probabilities that were oppositely yoked. For example, if the living category has a reward probability of 90%, the non-living category had a reward probability of 10%. Therefore, the subjects only had to learn the probability for one category, and simply assume the opposite probability for the other category.

The reward probability for a given image category remained stable until a change-point occurred, after which it changed to a random value between 0 and 1 (Figure 1d). Change-points occurred with a probability 0.16 on each trial. To facilitate learning, change-points did not occur in the first 20 trials of the task and the first 15 trials following a change-point. Each subject completed a unique task with pseudorandomized order of images that followed these constraints.

The objective was to maximize the total number of points earned. Subjects were advised to pay close attention to the value, probability, and category of each trial in order to decide whether it is better to PLAY or PASS. Subjects were thoroughly informed about the possibility of change-points, and that the two categories were oppositely yoked. They underwent a practice learning task in which the reward probabilities for the two categories were 1 and 0 to clearly demonstrate these features of the task. Subjects were awarded a bonus compensation proportional to the total points earned during the learning and memory tasks.

#### Memory task

During the memory task, subjects viewed 160 “old” images from the learning task intermixed with 160 “new” images (Figure 1c). Images were selected such that there was a new image for each unique exemplar from the learning task. This was to ensure that subjects had to make judgments about the actual image itself, rather than the fact that they saw a particular exemplar (e.g. “I remember seeing THIS desk” vs. “I remember seeing A desk”).

The order of old and new images was pseudorandomized. On each trial, a single image was presented, and the subject selected between OLD and NEW by pressing 1 or 0 on the keyboard, respectively (Figure 1c). Afterwards, they were asked to rate their confidence in the choice from 1 (Guess) to 4 (Completely certain). Subjects were not provided with correct/incorrect feedback on their choices.

#### Bayesian optimal learning model

The optimal learning model computed inferences over the probability of a binary outcome that evolves according to a change-point process. The model was given information about the true probability of a change-point occurring on each trial (H; hazard rate) by dividing the number of change-points by the total number of trials for each subject. For each trial, a change-point was sampled according to a Bernoulli distribution using the true hazard rate (CP ~ B(H)). If a change-point did not occur (CP = 0), the predicted reward rate (μ_t_) was updated from the previous trial (μ_t−1_). When a change-point did occur (CP = 1), μ_t_ ^was^ sampled from a uniform distribution between 0 and 1. The posterior probability of each trial’s reward rate given the previous outcomes can be formulated as follows:

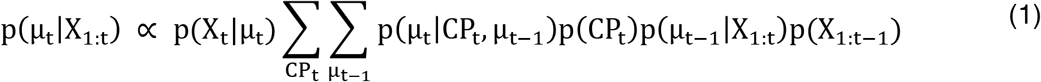

Where p(X_t_|μ_t_) is the likelihood of the outcomes given the predicted reward rate, p(μ_t_|CP_t_, μ_t−1_) represents the process of accounting for a possible change-point (when CP = 1, μ_t_ ~ U(0,1)), p(CP_t_) is the hazard rate, and p(μ_t−1_|X_1:t_) is the prior belief of the reward rate.

Using the model-derived reward rate, we quantified the extent to which each new outcome influenced the subsequent prediction as the learning rate in a delta-rule:

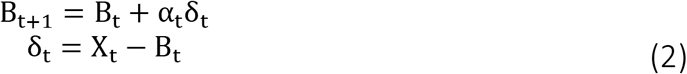

where B is the belief about the current reward rate, α is the learning rate, and δ is the prediction error, defined as the difference between the observed (X) and predicted (B) outcome. Rearranging, we were able to compute the trial by trial learning rate:

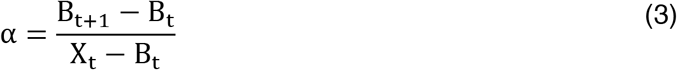

Trial by trial modulation of change-point probability (i.e. surprise) was calculated by marginalizing over μ_t_:

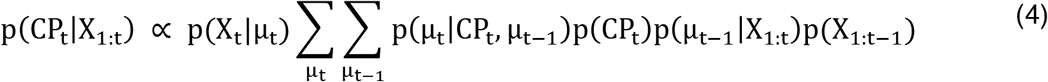

Uncertainty was determined by calculating the entropy of a discrete random variable X (i.e. reward rate) with possible values {x_1_,x_2_,…x_i_} for a finite sample (Shannon, 2001):

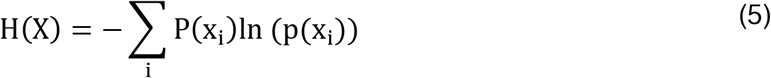

#### Descriptive analysis

Memory scores for each image were computed by transforming the recognition and confidence reports provided by the subject. On each trial of the recognition memory task, subjects first chose between “old” and “new”, then reported their confidence in that choice on a scale of 1-4. We converted these responses so that choosing “old” with the highest confidence (4) was a score of 8, while choosing “new” with the highest confidence was a score of 1. Similarly, choosing “old” with the lowest confidence (1) was a score of 5, while choosing “new” with the lowest confidence was a score of 4. As such, memory scores reflected a confidence-weighted measure of memory strength ranging from 1 to 8. These memory scores were used for all analyses involving recognition memory.

Statistical analyses were performed in a between-subject manner. For each subject, we computed the mean memory score of each trial type in question, then subtracted the overall average memory score of the subject. Therefore, the memory scores used in our analyses reflect the degree to which a certain trial condition led to better or worse subsequent memory compared to average performance within each subject.

Relationships between computational factors and memory scores were assessed by estimating the slope of the relationship between each computational factor and the subsequent memory score separately for each computational variable and subject. Statistical testing was performed using one sample t-tests on the regression coefficients across subjects (for overall effects) and two sample t-tests for differences between delay conditions (for delay effects). Regression coefficients (slopes) for individual subjects were related to individual differences in decision making task performance (coefficients from a GLM describing subject choices (play/pass) in terms of reward probability and trial value) using spearman rank order correlation.

To generate the descriptive figures, we performed a binning procedure for each subject to ensure that each point on the x axis contained an equal number of elements. For each subject, we divided the y variable in question into quartiles and used the mean y value of each quartile as the binned value. To plot data from all subjects on the same x axis, we first determined the median x value for each bin per subject, then took the average of the four bin median values across subjects. For figures containing more than one plot, we shifted the x values of each plot slightly off-center to avoid overlap of points.

A set of reinforcement learning models were fit directly to the subject behavior using a constrained search algorithm (fmincon in Matlab) that maximized the total log posterior probability of betting behavior given the optimal reward probability estimates, trial values, and prediction errors (Fig c&d). All models contained four fixed parameters that affected choice behavior: 1) a temperature parameter of the softmax function used to convert trial expected values into action probabilities, 2) a value exponent term that scales the relative importance of the trial value in making choices, 3) a play bias term to indicate a tendency to attribute higher value to gambling behavior, and 4) an intercept term for the effect of learning rate on choice behavior. The value estimated from gambling on a given trial was given by:

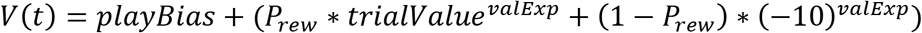

Where playBias is the play bias term, valExp is the value exponent, and P_rew_ is the reward probability inferred from the optimal model. The model fit with the above fixed parameters (the base model) was then compared to models that contained additional parameters that may affect trial-to-trial modulation of learning rate, including surprise, uncertainty, learning rate computed from the optimal Bayesian model, and subject choice behavior (play versus pass). In particular, learning rates were controlled through a logistic function of a weighted predictor matrix that included an intercept term as well as additional terms to capture the degree to which learning changed according to other factors. Maximum likelihood weights for each predictor in the matrix (as listed above) were estimated using gradient decent (fmincon in matlab) simultaneous with estimating the decision related parameters described above. The best fitting model was determined by computing the Bayesian information criterion (BIC) for each model, then comparing these values to that of the base model ^57^. Weak priors favoring normative learning parameters were used to regularize parameter estimates for parameter estimation but not model selection.

To compare subject behavior to model-predicted behavior, we simulated choice behavior using the model with the lowest BIC, which incorporated surprise and uncertainty variables in determining learning rate (Fig 2b). On each trial, we used the expected trial value (V(t)) computed above, and the parameter estimates of the temperature variable as inputs to a softmax function to generate choices.

#### Hierarchical regression model

Subject memory scores were modeled using a hierarchical mixture model that assumed that the memory score reported for each item and subject would reflect a linear combination of subject level predictors and item level memorability (Figure 6A). The hierarchical model was specified in STAN (http://mc-stan.org) using the matlabSTAN interface (http://mc-stan.org) ^58^. In short, memory scores on each trial were assumed to be normally distributed with a variance that was fixed across all trials for a given subject. The mean of the memory score distribution on a given trial depended on 1) a trial-to-trial task predictors that were weighted according to coefficients estimated at the subject level and 2) item-to-item predictors that were weighted by coefficients estimated across all subjects. Subject coefficients for each trial-to-trial task predictor were assumed to be drawn from a group distribution with a mean and variance offset by a delay variable, which allowed the model to capture differences in coefficient values for the two different delay conditions. All model coefficients were assumed to be drawn from prior distributions and for all coefficients other than the intercept (which captured overall memory scores) prior distributions were centered on zero. The code used to specify the hierarchical model is included as supplementary code.

### Experiment 2

#### Experimental procedure

In experiment 2, the learning task was modified to dissociate reward rate from reward prediction error. The reward probability of the two image categories (living vs. nonliving) were independent and set to either 0.8 or 0.2, allowing for a 2×2 design (0.8/0.8, 0.8/0.2, 0. 2/0.8, 0.2/0.2; Figure 7a). Change-points occurred with a probability 0.05 on every trial for the two categories independently. Change points did not occur for the first 20 trials of the task and the first 20 trials following a change point. Tasks were generated to contain at least one block of each trial type in the 2×2 design. Each subject completed a unique task with pseudorandomized order of images that followed these constraints. The task instructions explicitly stated that the two image categories had independent reward probabilities that need to be tracked separately. The rest of the task, including the recognition memory portion, was identical to that of Experiment 1.

#### Subjects

A total of 279 subjects (157, no delay condition; 122, 24hr-delay condition) completed the task via Amazon Mechanical Turk. 105 subjects (64, no delay; 41, 24hr-delay) were excluded from analysis because they previously completed a prior version of the task or didn’t meet our criteria of above-chance performance in the learning task. Therefore, the final sample had a total of 174 subjects (93, no delay, 81, 24hr-delay; 101 males, 71 females, 2 no response) with the age of 34.0 ± 9.1 (mean ± SD). Informed consent was obtained in a manner approved by the Brown University Institutional Review Board.

## Data availability

The data from both experiments and the scripts used to analyze and model the data are available from the authors upon request.

## Author Contributions

All authors designed the experiment and wrote the manuscript. AIJ collected the data and MRN developed the computational mdoels. MRN and AIJ designed and performed behavioral analysis.

## Competing Interests

None of the authors have any competing interests to report, financial or otherwise.

## Acknowledgements

We thank Anne Collins for helpful comments on the experimental design. This work was funded by NIH grants F32MH102009 and K99AG054732 (MRN), NIMH R01 MH080066-01 and NSF Proposal #1460604 (MJF), and R00MH094438 (DGD). The funders had no role in study design, data collection and analysis, decision to publish or preparation of the manuscript.

## Supplementary figures

**Supplementary figure 1.**
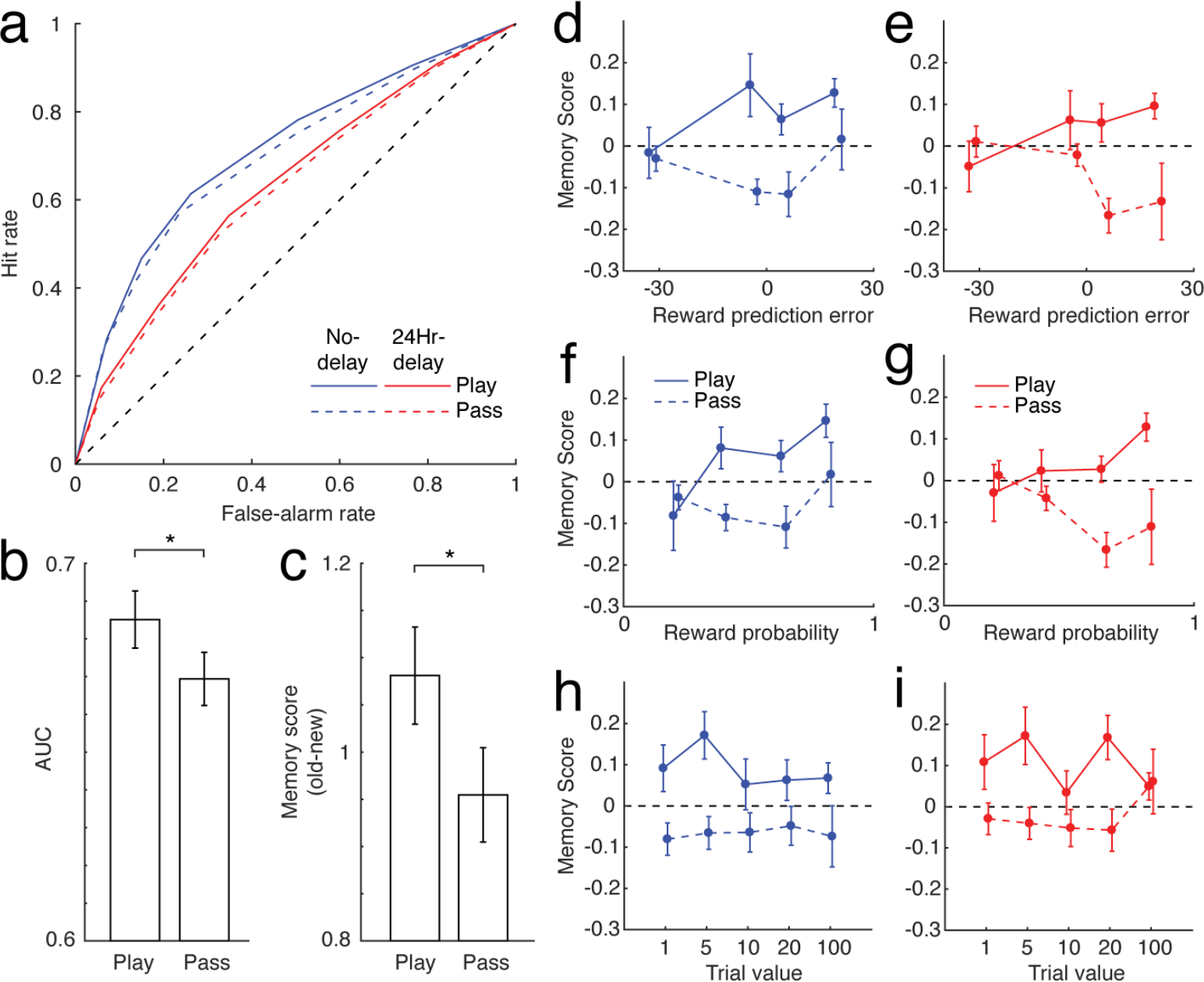
Replication of main effects after omitting all trials in which subsequent memory had low confidence (“guess”). To rule out the possibility that low confidence trials are driving our results or adding unnecessary noise, we repeated the main findings of the study after omitting all trials in which the confidence score was 1 (both target and foil). (a) The ROC curves for play/pass trials. (b) Area under the ROC curve was greater for play versus pass (t(198) = 2.78, p = 0.0060; group difference, t(197) = −0.083, p = 0.93). (c) Mean pairwise difference in memory score between the “old” images and their semantically-matched foil images was greater for play versus pass (t(197) = 3.49, p < 0.001; group difference, t(196) = 89, p = 0.37). (d,e) Positive relationship between imRPE and subsequent memory (t(198)=2.48, p=0.014; group difference, t(197)=−0.33, p=0.74). (f,g) Positive relationship between reward probability and subsequent memory (t(198)=2.99, p=0.0031, group difference, t(197)=0.078, p=0.94). (h,i) No relation between reward value and subsequent memory (t(198)=−1.74, p=0.084, group difference, t(197)=0.82, p=0.41).

**Supplementary figure 2:**
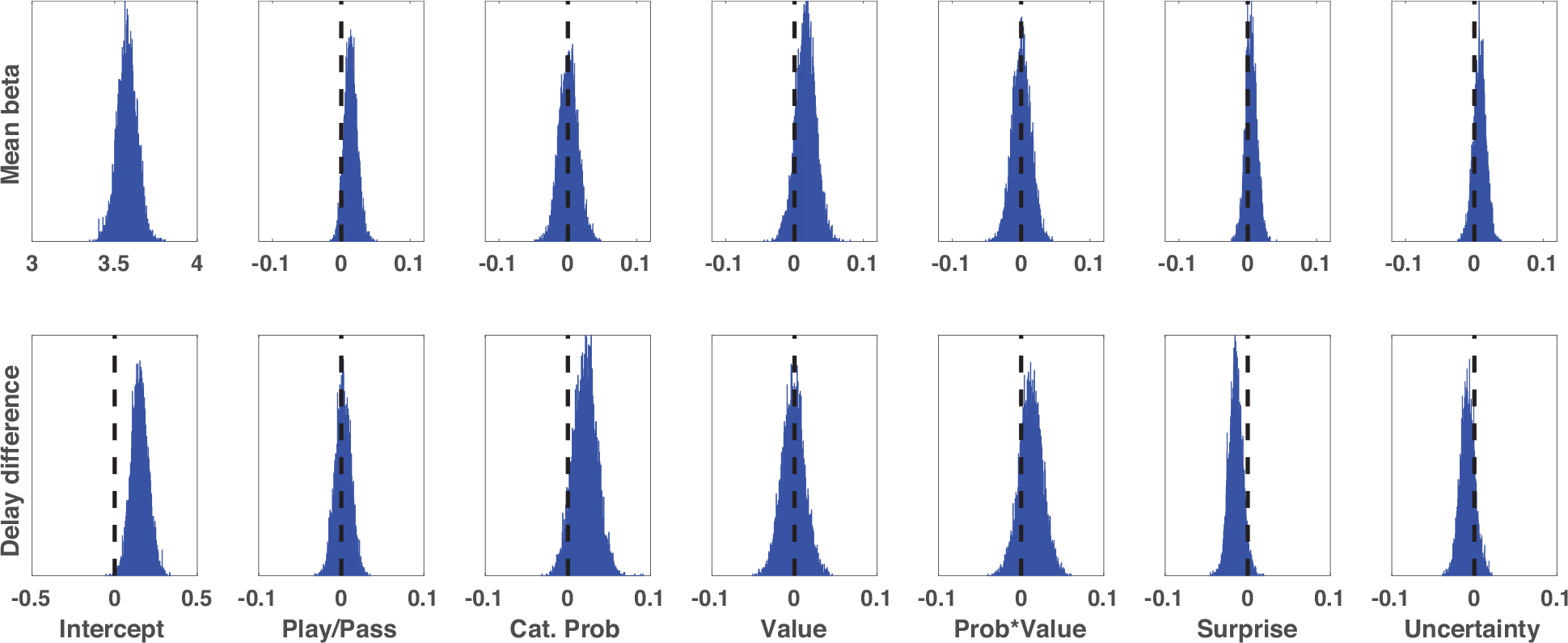
Hierarchical regression estimating memory scores for foil items. Posterior probability densities for mean predictor coefficients (μ_x_; top row) and delay condition parameter difference (*D*_x_; bottom row) estimated through MCMC sampling over the graphical model described in figure 6A informed by data for foil items semantically matched to the images presented in the decision making task. Unlike fits to the target items (Figure 6B), coefficients for task predictors related to reward probability and play/pass did not deviate appreciably from zero. However, delay condition difference coefficients were positive for the intercept term, indicating that subjects in the 24 hour delay condition tended to report higher memory scores for foils, consistent with the poorer discriminability in the 24 hour delay condition (see figure 3C).

**Supplementary figure 3.**
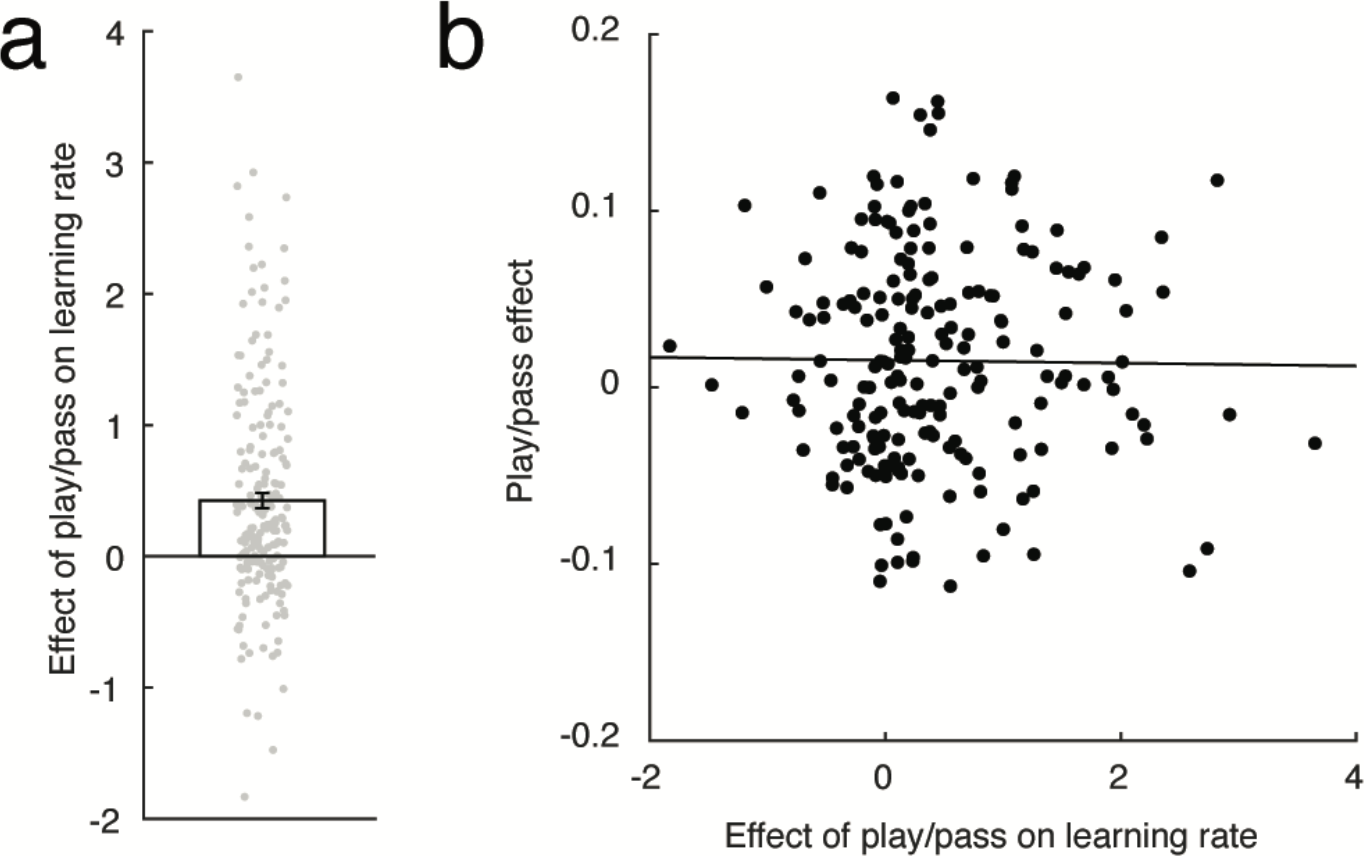
Subject learning rates differed across choice conditions but this difference did not predict the strength of choice effects on subsequent memory. **a:** Parameter estimates quantifying the degree to which learning rate is modulated by subject play/pass decisions in the choice model (positive values indicate more learning on after play decisions) for all subjects in experiment 1 (gray points) and the mean/SEM values across subjects (bar/error line; mean parameter estimate = 0.42, t = 7.18, p < 0.001). **b:** The degree to which subjects (black points) adjusted learning rate according to play/pass behavior (abscissa) showed no association with the degree to which they enhanced memory of items presented on play trials (ordinate; spearman rho = 0.045, p = 0.53).

**Supplementary figure 4.**
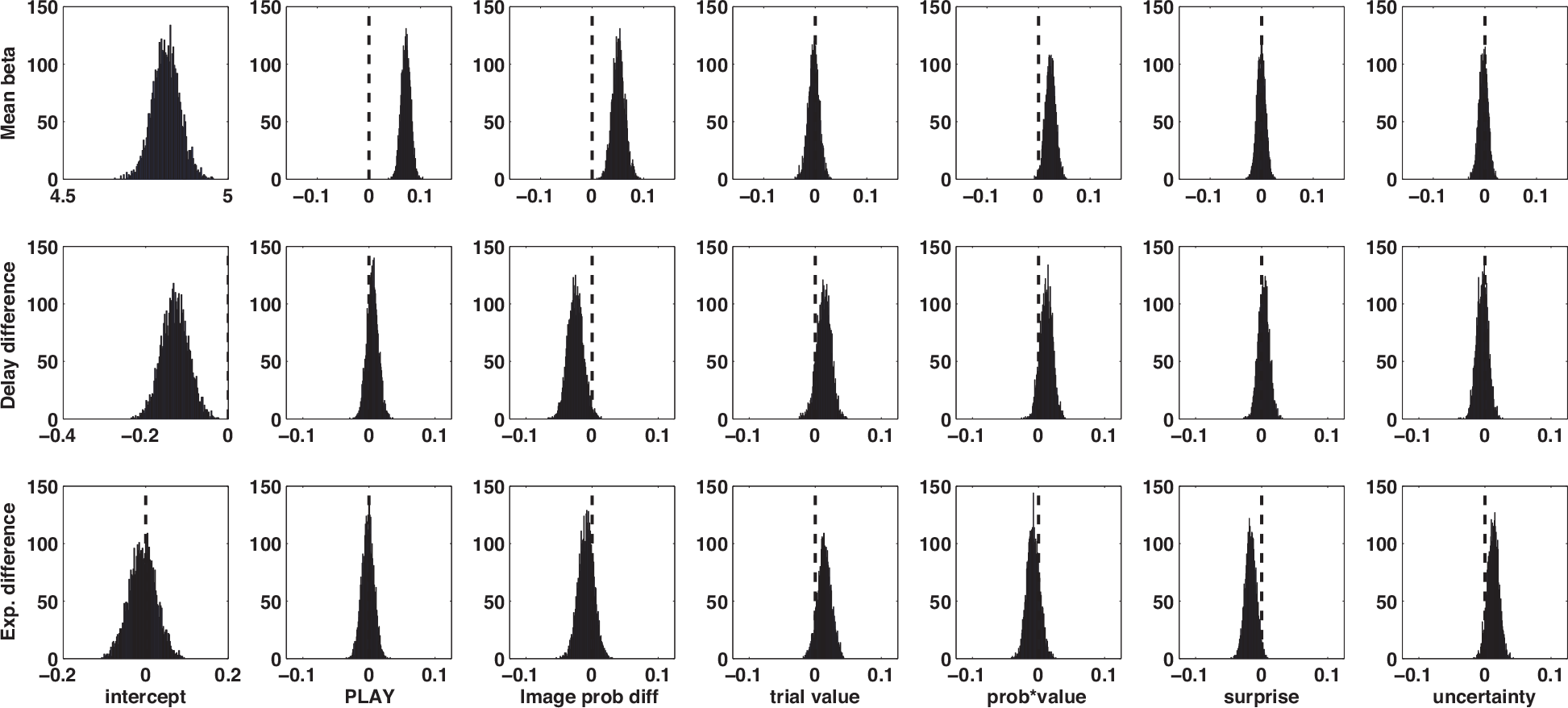
Coefficient estimates for hierarchical model fit to all subjects across both experiments. **Top:** Posterior probability density over mean coefficient estimates at the population level for each parameter in the hierarchical model fit to subject memory scores. Leftmost column reflects the intercept indicating overall memory scores for old items, whereas all other columns reflect the degree to which learning task-related factors affected subsequent memory. The model included two separate terms to model the probability associated with the shown image category (“cat prob(play)”; third column) and the probability associated with the non-displayed image category (“non-cat prob(play”; fourth column) despite the fact that these two terms were perfectly anti-correlated for all participants who completed experiment 1. **Middle:** Posterior probability density over delay difference estimates for each parameter in the hierarchical model. **Bottom:** Posterior probability density over experiment difference estimates quantifying the difference in coefficient values across the two experiments for each parameter in the hierarchical model.

**Supplementary code:**
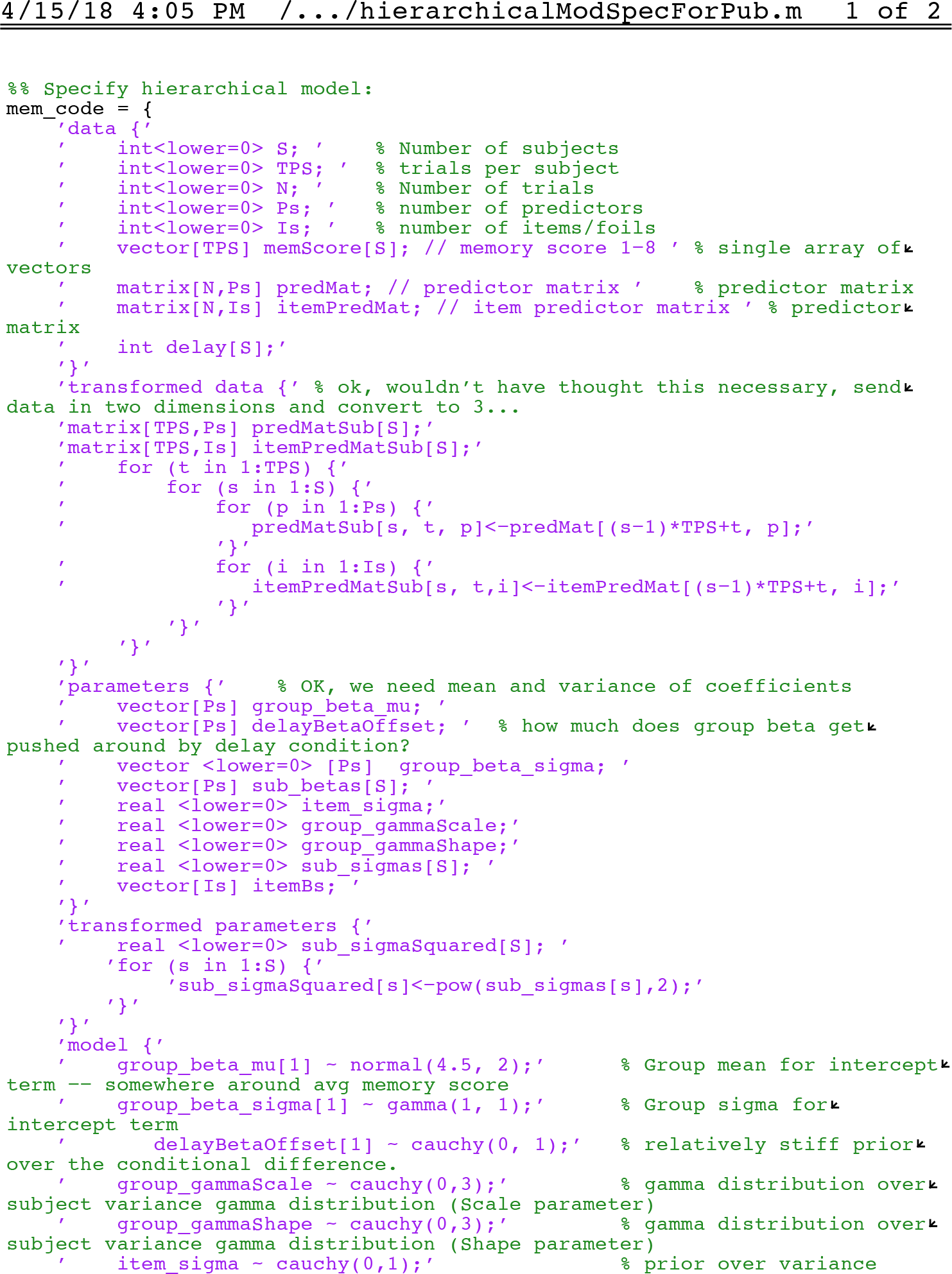
Hierarchical regression model for experiment 1 spefified in matlabStan. All code and data will be available by the authors upon request.

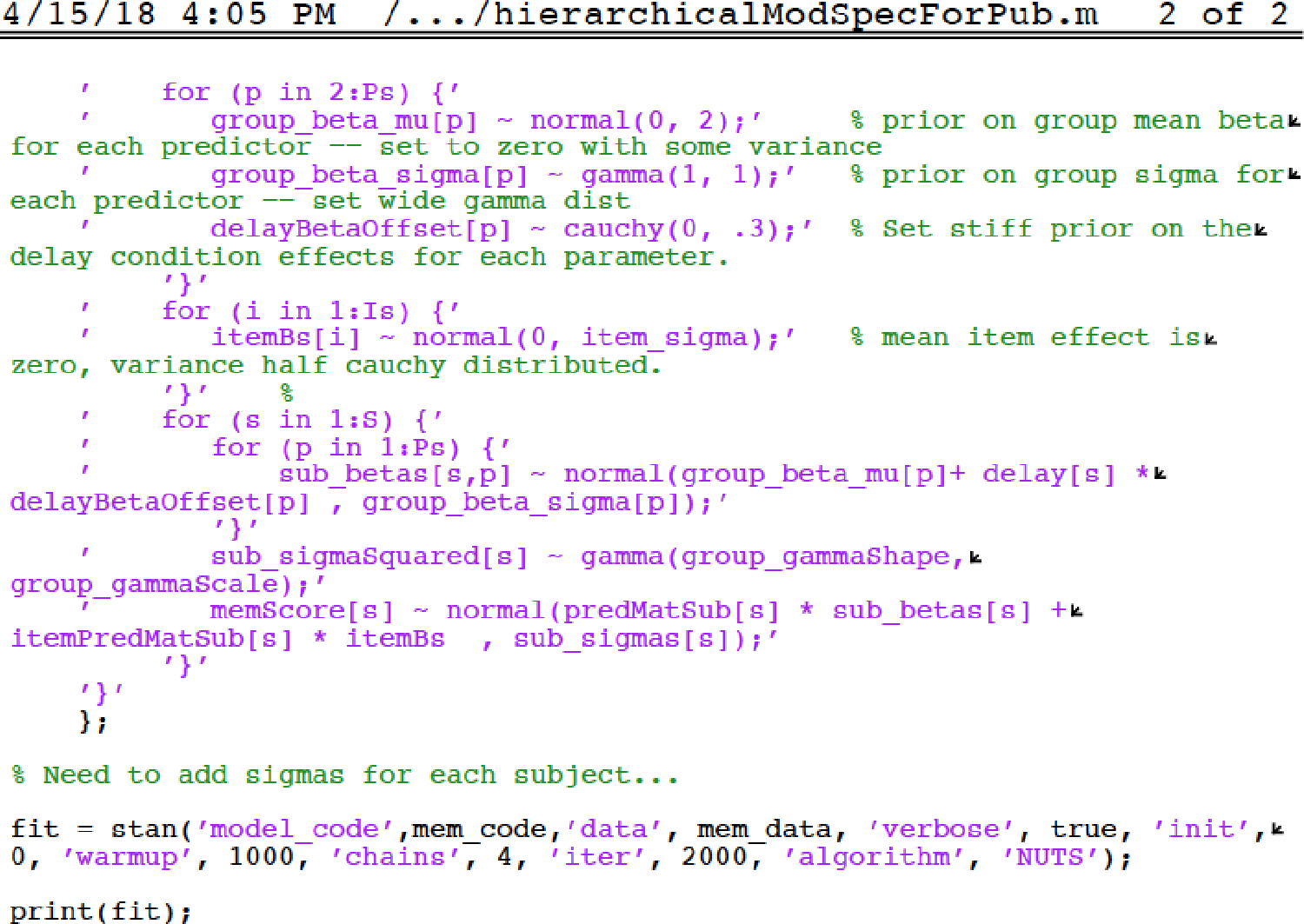

